# SillyPutty: Improved clustering by optimizing the silhouette width

**DOI:** 10.1101/2023.11.07.566055

**Authors:** Polina Bombina, Dwayne Tally, Zachary B. Abrams, Kevin R. Coombes

## Abstract

Unsupervised clustering is an important task in biomedical science. We developed a new clustering method, called SillyPutty, for unsupervised clustering. As test data, we generated a series of datasets using the Umpire R package. Using these datasets, we compared SillyPutty to several existing algorithms using multiple metrics (Silhouette Width, Adjusted Rand Index, Entropy, Normalized Within-group Sum of Square errors, and Perfect Classification Count). Our findings revealed that SillyPutty is a valid standalone clustering method, comparable in accuracy to the best existing methods. We also found that the combination of hierarchical clustering followed by SillyPutty has the best overall performance in terms of both accuracy and speed.

**Availability:** The SillyPutty R package has been submitted to the Comprehensive R Archive Network (CRAN). Code to perform and analyze the simulations described here can be found in a Git project hosted at https://gitlab.com/krcoombes/sillyputty.

## 1 Introduction

In modern data analysis, uncovering inherent groupings within a collection of patterns, data points, or objects has significant value. Unsupervised clustering is a crucial tool to tackle this challenge. It is widely acknowledged that there is no one-size-fits-all approach [1,2]. It is often important to tailor the cluster analysis method to the specific requirements of the data at hand, based on domain expertise and the ultimate purpose of clustering [3].

Cluster algorithms have been used, and new ones proposed, for decades. Ward’s linkage rule for hierarchical clustering was proposed in 1963 [4]. The popular K-means method was introduced by 1967 [5]. The methods of partitioning around medoids (PAM) and clustering large applications (CLARA) were introduced by Kaufman and Rooseeuw in the late 1980’s [6,7]. After the development of high throughput biological assays (starting with gene expression microarrays) in the late 1990’s, the following decade saw the introduction of new algorithms designed to work with larger data sets. These included spectral clustering from 2001 [8–10], and subspace clustering [11,12], which started with the CLIQUE algorithm in 1998 [13].

Different algorithms rely on different models of the shapes of clusters, and they optimize different metrics. K-means, for example, is a heuristic algorithm to minimize the within-group sum-of-squares. PAM is a related method that generalizes the K-means optimization target, the Euclidean distance, to arbitrary distance metrics. Existing clustering algorithms have used other measures to guide their heuristics, including density-based methods [14] or gradient descent optimization [15].

In 1987, Rousseeuw introduced an idea known as the silhouette width (SW) [16]. The SW is defined for each clustered object, producing a value in the interval [−1, +1]. The SW gives us an idea of how strongly we should believe that an object has been assigned to the correct cluster. Positive values indicate good clustering, negative values indicate that the object is closer to a different cluster. To date, its main application has been to use the mean silhouette width (ASW) over all objects to assess the overall quality of a clustering assignment and to decide on the “correct” number of clusters [7].

Inspired by the applications of ASW to assess cluster quality, we had the idea that a heuristic clustering method based on maximizing the mean silhouette width might be useful. We developed a novel clustering method, called “SillyPutty”, that relies on the individual SW values to drive the heuristic. To our knowledge, no one has previously used the silhouette width of individual objects in this way.

In this manuscript, we present a series of simulations to compare our new SillyPutty algorithm to the top performing methods arising from a recent study that compared existing clustering algorithms [17]. We use both internal and external clustering validation metrics to evaluate the algorithms [18]. In the “Materials and methods” section, we review the silhouette width and describe the SillyPutty algorithm in detail. Next, we outline the data generation methodology and describe the performance metrics used to compare the algorithms. Finally, we present and analyze the performance results.

## 2 Materials and methods

All computations were performed using R version 4.3.1 (2023-06-16 ucrt) of the R Statistical Programming Environment [19].

### 2.1 Existing cluster algorithms

The recent study conducted by Rodriguez and colleagues [17] involved a thorough evaluation wherein they compared nine frequently employed clustering methods available in the R language [19]. In our work, for each simulated data set, we compare the performance of SillyPutty with the “winners” of their analysis: PAM, CLARA, Hierarchical, K-means, Spectral, and Subspace.

We used the implementations of both hierarchical clustering (hclust) with Ward’s linkage [4] and the K-means (`kmeans`) algorithm [5] from version 4.3.1 of the stats R package. We used the implementations of the PAM (`pam`; [6]) and CLARA (clara; [7]) algorithms from version 2.1.4 of the cluster R package. We used the implementation of spectral clustering (`specc`; [8,10]) from version 0.9.32 of the kernlab R package. Finally, we used the implementation of subspace clustering (`hddc`; [11]) from version 2.2.0 of the `HDclassif` R package.

### 2.2 Silhouette width

The silhouette width is a well-known and popular measure of how well each data point fits its designated cluster. Let *X* = {*x*_1_, …, *x*_*n*_} ∈ *R*^*F*^ be a finite point set along with a pairwise distance metric *D*. Here *F* is the number of features measured at each data point. Define *K* to be the (true) number of clusters. The silhouette width for an observation *x*_*i*_ ∈ *X* is calculated as follows:

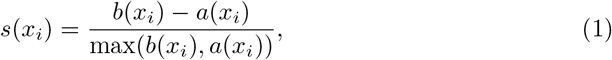

Here *b*(*x*_*i*_) is a measure of separation; that is, the minimum (over other clusters) average distance between *x*_*i*_ and the data points in the other cluster. Similarly, *a*(*x*_*i*_) is a measure of cohesion; that is, the average distance between *x*_*i*_ and every data point in its assigned cluster.

The silhouette width *s*(*x*_*i*_) falls within the range [−1, 1]. A value near 1 signifies that the data point is well-clustered, indicating distinct separations between clusters and strong cohesion within each group. On the other hand, a value closer to 0 implies overlapping clusters, while a value near −1 suggests that the data point has been miscassified.

### 2.3 SillyPutty algorithm

We propose a novel clustering algorithm, called SillyPutty, in this section. SillyPutty is a heuristic algorithm based on the concept of silhouette widths. Its goal is to iteratively optimize the cluster assignments to maximize the average silhouette width. SillyPutty starts with any given set of cluster assignments, either user-specified, randomly chosen, or obtained from other clustering methods. SillyPutty enters a loop where it iteratively refines the clustering. The algorithm calculates the silhouette widths for the current clustering. Then it identifies the data point with the lowest (most negative) silhouette width, which implies a potential misclassification. The algorithm reassigns this data point to the cluster to which it is closest. The loop continues until all data points have non-negative silhouette widths, or an early termination condition is reached. SillyPutty halts under three conditions:

- When all data points are well-classified, as evidenced by having all silhouette width non-negative, indicating convergence.
- When the maximum number of iterations is reached, to avoid infinite loops.
- When it detects a possible infinite loop by finding the same vector of silhouette widths within the last *n* iterations, where *n* is user-specified.

Upon convergence or termination, SillyPutty returns the refined clustering, along with the corresponding silhouette widths and other relevant information. A summary of the SillyPutty workflow is shown in Fig 1.

**Fig 1.**
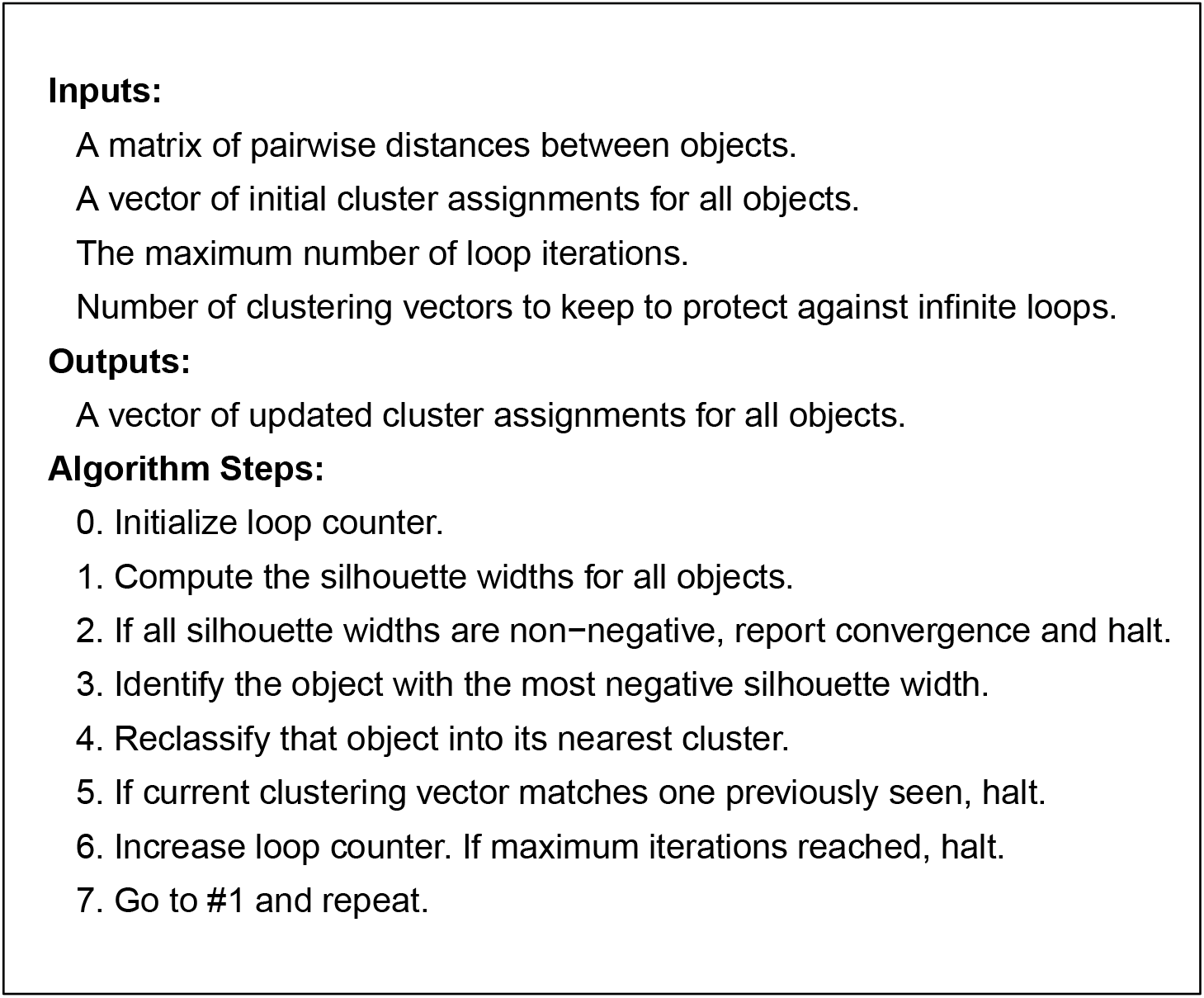
SillyPutty algorithm.

### 2.4 Axiomatic characterization of the ASW

Since the SillyPutty algorithm is rooted in the ASW concept, it is useful to verify how well ASW, a metric for cluster quality, aligns with theoretical principles. Batool and Hennig [20] used an axiomatic approach to assess ASW. They found that, as a clustering quality measure,

- ASW is scale invariant; that is, the result remains the same even when all dissimilarities are uniformly scaled by a constant factor.
- ASW is consistent; that is, reducing or maintaining all within-cluster dissimilarities while increasing or maintaining all between-cluster dissimilarities, does not alter the result.
- ASW is rich; that is, every conceivable clustering can be generated by an appropriate dissimilarity measure

### 2.5 Hybrid approaches

The standalone `SillyPutty` algorithm starts with purely random cluster assignments, repeating the algorithm with (by default) 100 different random starting points. Since `SillyPutty` can start with any cluster assignments, we also investigated the efficacy of following other clustering methods by applying `SillyPutty` to their cluster assignments. As `SillyPutty` requires a distance matrix as one of its inputs, we compute distances using the Euclidean metric. Our goal with this part of the study was to assess whether integrating `SillyPutty` with existing clustering techniques can improve their performance.

### 2.6 Simulated datasets

To generate realistic datasets, we employed version 2.0.10 of the `Umpire` (Ultimate Microarray Prediction, Inference, and Reality Engine) R package [21,22], which enables us to generate gene-expression-like data on the log-transformed scale from multiple clusters, with known “ground truth”. These data mimic the characteristics of actual cancer transcriptomics data.

We generated data sets that differ in the number of clusters (3, 6 or 12), the sample size (600 or 1000), and the feature size, (e.g., number of genes; 5000 or 10000). Moreover, we modify the data by applying different levels of noise (additive on the log scale), categorized as low, medium, or high. Mathematically, if *Y*_*gi*_ represents the observed expression of gene *g* in sample *i, S*_*gi*_ represents the true biological signal of gene *g* in sample *i*, and *ϵ*_*gi*_ defines the additive noise for gene *g* in sample *i*, then the noise model can be represented as:

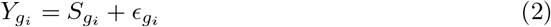

In our framework, *ϵ* ∼ *N* (*ν, τ*). We setν = 0.1 and vary *τ*, using a gamma distribution with different means to model differing standard deviations *τ* .

The summary of distinct datasets that were generated is shown in Table 1. We repeat each set of parameters 19 times, so that our study encompasses a total of 513 individual simulations. To illustrate the simulated data, we present graphical summaries of four data sets from one replicate run (Fig 2). These include (A) a data set that should be easy to classify, with three clusters, 600 samples, 5000 features, and low noise; (B) another “easy” data set, with six clusters, 1000 samples, 5000 features, and low noise, (C) a data set of “intermediate” difficulty, with six clusters, 600 samples, 10000 features, and medium noise; and (D) a “hard” data set, with twelve clusters, 1000 samples, 10000 features, and high noise.

**Table 1.** List of cancer genomics datasets generated by the Umpire package.

| Dataset | Clusters | Noise | Samples | Features |
| --- | --- | --- | --- | --- |
| 1 | 12 | High | 1000 | 10000 |
| 2 | 12 | Low | 1000 | 10000 |
| 3 | 12 | Low | 1000 | 5000 |
| 4 | 12 | Low | 600 | 10000 |
| 5 | 12 | Low | 600 | 5000 |
| 6 | 12 | Medium | 1000 | 10000 |
| 7 | 12 | Medium | 1000 | 5000 |
| 8 | 12 | Medium | 600 | 10000 |
| 9 | 12 | Medium | 600 | 5000 |
| 10 | 3 | High | 1000 | 10000 |
| 11 | 3 | Low | 1000 | 10000 |
| 12 | 3 | Low | 1000 | 5000 |
| 13 | 3 | Low | 600 | 10000 |
| 14 | 3 | Low | 600 | 5000 |
| 15 | 3 | Medium | 1000 | 10000 |
| 16 | 3 | Medium | 1000 | 5000 |
| 17 | 3 | Medium | 600 | 10000 |
| 18 | 3 | Medium | 600 | 5000 |
| 19 | 6 | High | 1000 | 10000 |
| 20 | 6 | Low | 1000 | 10000 |
| 21 | 6 | Low | 1000 | 5000 |
| 22 | 6 | Low | 600 | 10000 |
| 23 | 6 | Low | 600 | 5000 |
| 24 | 6 | Medium | 1000 | 10000 |
| 25 | 6 | Medium | 1000 | 5000 |
| 26 | 6 | Medium | 600 | 10000 |
| 27 | 6 | Medium | 600 | 5000 |

**Fig 2.**
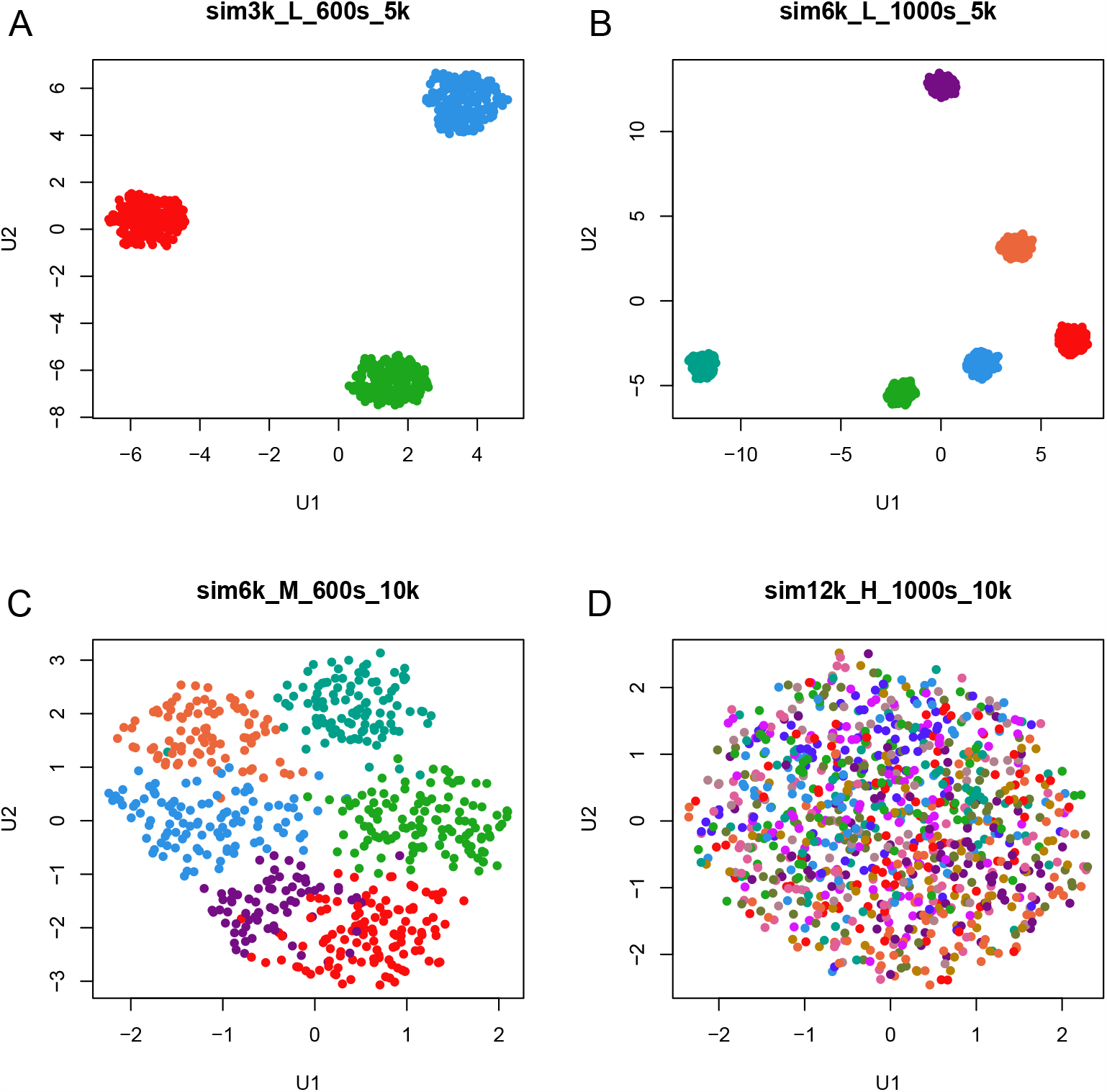
UMAP plots of four examples of simulated data sets using different parameters. (A) 3 clusters, 600 samples, 5000 features, low noise. (B) 6 clusters, 1000 samples, 5000 features, low noise. (C) 6 clusters, 600 samples, 10000 features, medium noise. (D) 12 clusters, 1000 samples, 10000 features, high noise.

### 2.7 Comparative evaluation

For each set of simulation parameters, we computed the average performance across the repetitions to determine the most effective algorithm. Our evaluation criteria encompass both external measures (the Adjusted Rand Index (ARI) and Entropy), and internal measures (ASW and the “Normalized Within-group Sum-of-Squares” (NWSS)). Finally, we consider a measure we call the Perfect Classification Count (PCC). Furthermore, we assess and compare the computational runtime of all algorithms.

#### 2.7.1 Adjusted Rand index (ARI)

The ARI is a measure of agreement between two sets of clustering assignments. It is computed as follows:

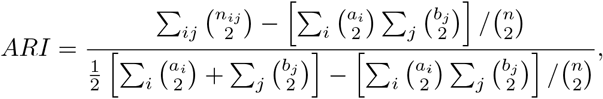

where *n*_*ij*_ is the number of data points that are in the same cluster in both clustering solutions, *a*_*i*_ is the total number of data points in cluster *i* in the first clustering solution, *b*_*j*_ is the total number of data points in cluster *j* in the second clustering solution, and *n* is the total number of data points.

The “external” aspect of ARI arises because we compare any algorithm-derived cluster to the known truth. It is the metric of choice for assessing overall agreement between clustering methods, while taking into account the potential occurrence of coincidental agreement [18].

#### 2.7.2 Entropy

Entropy measures the degree of impurity or disorder within each cluster. The entropy of a single cluster *S*_*i*_ is defined to be

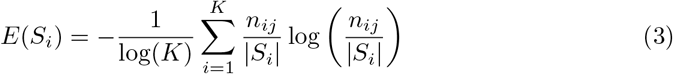

Here, *n*_*ij*_ is the number of data points in the same cluster in both clustering solutions for cluster *S*_*i*_, |*S*_*i*_| is the total number of data points in cluster in the second clustering solution, *K* is the number of clusters.

Global entropy is defined to be the sum of the individual cluster entropies, weighted by the cluster size:

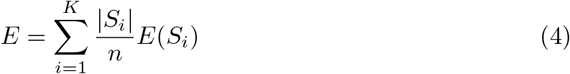

Global entropy takes values between 0 and 1. Lower entropy values indicate that the cluster assignments are more homogeneous which one expects will better correspond to the ground truth. To calculate the entropy, we call the `external validation` function from version 1.3.1 of the ClusterR R package. The entropy measure is also “external”, since we compare each clustering solution to the known ground truth.

#### 2.7.3 Mean silhouette width

The SW formula was presented above (Eqn (1)). We use the `silhouette` function from version 2.1.4 of the `cluster` R package to calculate the ASW.

#### 2.7.4 Normalized within-group sum-of-squares

The within-group sum of square errors is equivalent to the sum of distances of each point from the centroid of its assigned cluster. This can be written as:

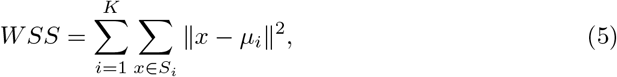

where *μ*_*i*_ is the centroid of the cluster *S*_*i*_. This equation is the same as the K-means objective function. For our analysis, we wrote our own function wss to return the WSS for a (numeric) data frame. Then we normalize WSS by dividing by the WSS from the true cluster assignments so that the result does not depend on the size of the data set. We expect NWSS to take values in [1, ∞).

#### 2.7.5 Perfect classification count

We introduce a metric called Perfect Classification Count (PCC). PCC is the number of times (out of all 513 simulations) that a method correctly classifies all samples in a simulated data set. The percentage of perfect classifications is an empirical estimate of the probability that a method will produce perfect (“true”) clusters.

## 3 Results

We recorded the average run time for each method. The values we recorded are for running each method on a complete set of 27 different combinations of simulation parameters. The running times of algorithms are provided in Table 2. We note that the standalone `SillyPutty` clustering exhibits the slowest performance, with an average convergence time of 9629 seconds. Hierarchical clustering outperforms the rest of algorithms demonstrating the highest time efficiency than any other method.

**Table 2.** Average running time of algorithms across all simulations.

| Method | Running Time (seconds) |
| --- | --- |
| Hierarchical | 1.0 |
| Kmeans+Silly<sup>a</sup> | 4.7 |
| Sub<sup>b</sup>+Silly<sup>a</sup> | 5.5 |
| Hier<sup>c</sup>+Silly<sup>a</sup> | 16.9 |
| Spec<sup>c</sup>+Silly<sup>a</sup> | 26.1 |
| PAM+Silly<sup>a</sup> | 40.3 |
| CLARA+Silly<sup>a</sup> | 46.3 |
| Kmeans | 99.2 |
| CLARA | 120.8 |
| Spectral | 416.2 |
| Subspace | 3214.8 |
| PAM | 8925.7 |
| SillyPutty | 9628.9 |

<sup>a</sup> Hybrid methods report the time for SillyPutty.
<sup>b</sup> Sub = Subspace clustering.
<sup>c</sup> Hier = Hierarchical clustering.
<sup>d</sup> Spec = Spectral clustering.

In Fig 3, we show the distributions of values for four performance metrics for each method across all simulations. It is clear that the methods PAM and CLARA are far inferior to other methods, while Spectral and Hierarchical have intermediate performance. In general, following almost any method by applying `SillyPutty` makes the results substantially better.

**Fig 3.**
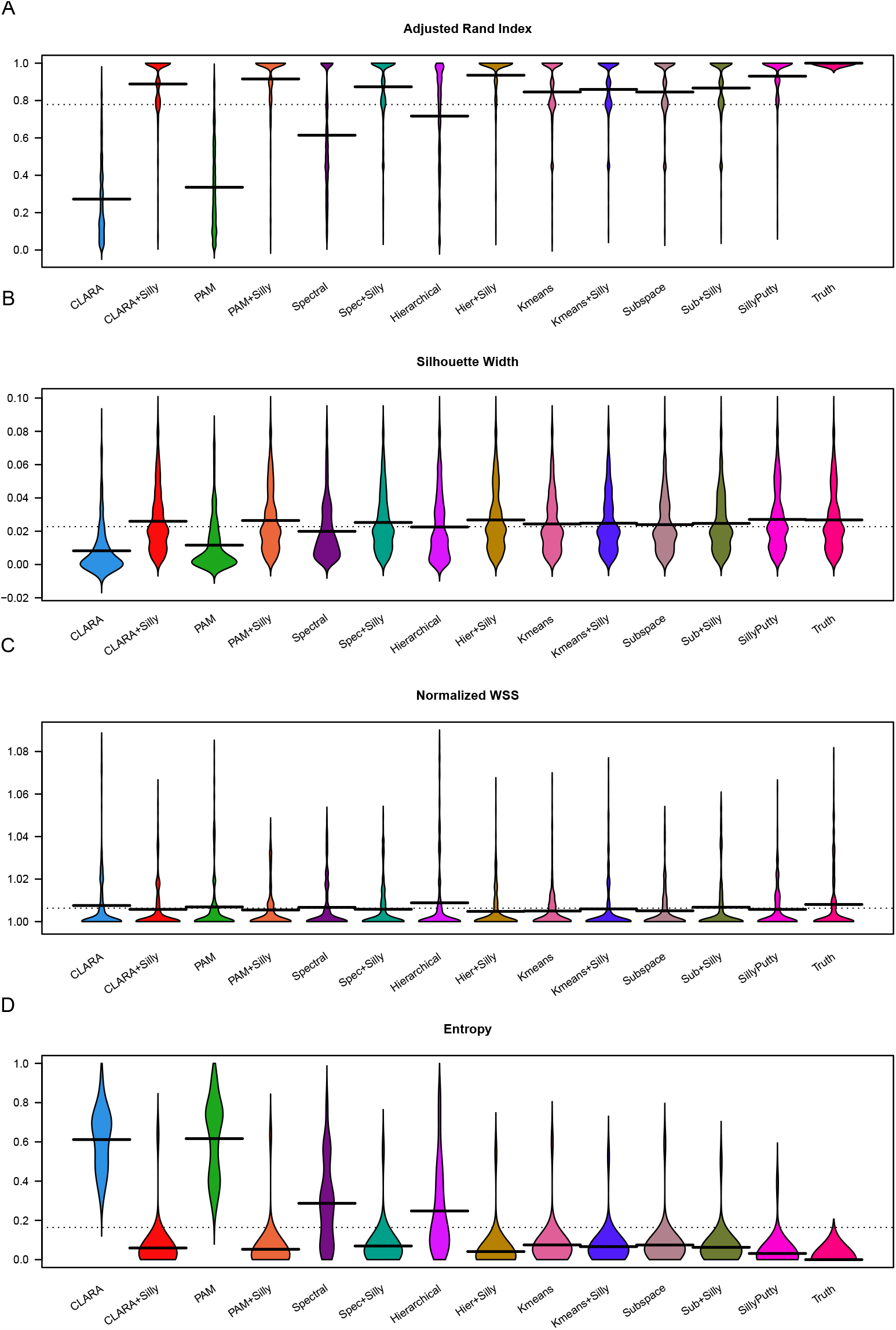
Evaluation of algorithms. Values over 19 replicate simulated sets were averaged. Distributions of averaged values over 27 sets of simulation parameters are displayed in ‘bean plots’ for each of the 13 methods (plus true clusters) (A) Adjusted Rand Index. (B) Mean silhouette width. (C) Normalized within-group sum of squares. (D) Entropy.

Moreover, in Table 3, we report a summary of mean values of each performance metric for each method across all simulations. By these measures, `SillyPutty` from random starts has the best performance of standalone algorithms. CLARA and PAM have the worst performance, with hierarchical clustering (HC) and Spectral clustering in the middle. Applying `SillyPutty` after running another algorithm makes very little change to K-means or Subspace clustering, but it dramatically improves the performance of the four inferior methods, causing all four to perform better than K-means or Subspace. The best overall performance in four out of five metrics is found by first doing hierarchical clustering followed by `SillyPutty` (HCSilly), with perfect performance in 70% of simulations, average ARI of 0.94, average SW of 0.027, and average NWSS of 1.

**Table 3.** The averaged values of the validity indices for all clustering methods across all simulated data experiments.

| Method | SW | ARI | NWSS | Entropy | PCC |
| --- | --- | --- | --- | --- | --- |
| CLARA | 0.008 | 0.272 | 1.008 | 0.612 | 0 |
| PAM | 0.012 | 0.336 | 1.007 | 0.617 | 0 |
| Spectral | 0.020 | 0.614 | 1.007 | 0.287 | 122 |
| Hierarchical | 0.023 | 0.717 | 1.009 | 0.248 | 13 |
| Subspace | 0.024 | 0.846 | 1.005 | 0.075 | 210 |
| Kmeans | 0.024 | 0.846 | 1.005 | 0.075 | 226 |
| Kmeans+Silly | 0.025 | 0.859 | 1.006 | 0.066 | 222 |
| Sub+Silly | 0.025 | 0.867 | 1.007 | 0.063 | 229 |
| Spec+Silly | 0.025 | 0.873 | 1.006 | 0.070 | 255 |
| CLARA+Silly | 0.026 | 0.888 | 1.006 | 0.059 | 275 |
| PAM+Silly | 0.026 | 0.915 | 1.005 | 0.053 | 322 |
| SillyPutty | 0.027 | 0.930 | 1.006 | 0.031 | 318 |
| Hier+Silly | 0.027 | 0.935 | 1.005 | 0.041 | 355 |
| Truth | 0.027 | 1.000 | 1.008 | 0.000 | 513 |

We next looked at the ways in which the simulation parameters affected the Adjusted Rand Index (Fig 4). We performed this analysis for the number of clusters, the number of samples, the number of features, and the noise level. It is clear that more clusters, more samples, more features, or more noise decrease the ARI. We also looked at combinations of parameters and concluded that the effects appeared to be independent and additive (see supplementary material). We performed similar analyses for other outcome measures (ASW, Entropy) and found qualitatively similar results. However, NWSS appeared not to be significantly impacted by the factors under consideration.

**Fig 4.**
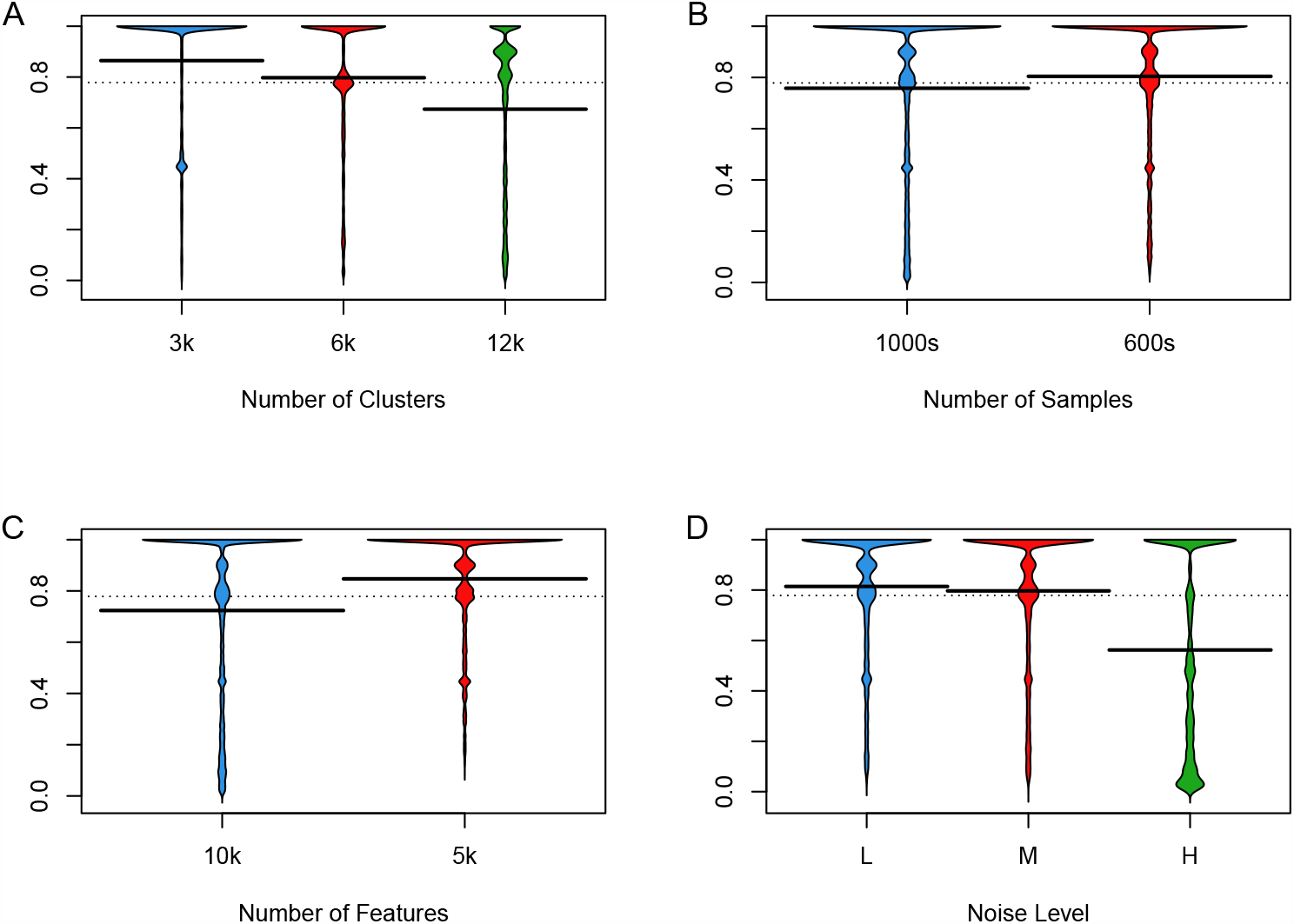
Effect of simulation parameters on Adjusted Rand Index. (A) Clusters. (B) Samples (C) Features. (D) Noise

## 4 Discussion

In this study, we conducted a comprehensive comparison of unsupervised clustering methods using simulated data. Our goal was to identify the top-performing method and assess the potential of SillyPutty, a novel clustering algorithm that optimizes silhouette width, in improving overall clustering performance.

In terms of accuracy, `SillyPutty` had better performance than any of the existing clustering methods. It achieved perfect agreement with the known ground truth in 318 (61.9%) out of 513 data sets, its mean ARI score was 0.930, and its mean entropy was 0.031. By all three measures, it was the best of the standalone algorithms.

However, this accuracy came at a cost; `SillyPutty` was by far the slowest standalone algorithm (taking an average of 9629 seconds, or 2.67 hours, over a 27-data-set simulation run). These large elapsed times appear to arise from two sources. First, we computed the length of time to run the base algorithm from 100 random starts. So, the mean time for a single run is only 96 seconds, about the same time taken by K-means. A second source of longer run times is the amount of work that must be performed when starting from random cluster assignments. The amount of time reported in Table 2 to run `SillyPutty` in “hybrid” methods (that is, after first running one of the other algorithms) was considerably shorter, rangeing from 4.7 seconds after K-means to 46.3 seconds after CLARA. It is interesting to note that those times appear to be related to the quality of the initial clustering. The better the initial clustering, the faster `SillyPutty` completes its task. The fastest combined mean run time (17.9 seconds) for a hybrid method was achieved by first performing hierarchical clustering (which is extremely fast) and then running SillyPutty.

We also found that the same hybrid combination, hierarchical clustering followed by SillyPutty, was the best method overall in our experiments. It had a perfect classification count of 355/513 (69.2%), a mean ARI of 0.935, and a mean entropy of 0.041. These results are significantly better than the best existing standalone method, K-means, with PCC = 236/513 (46.0%), ARI = 0.846, and entropy = 0.075. In fact, following any existing method with `SillyPutty` improved the performance. Of course, bigger improvements were observed when following algorithms that had worse performance to start with (witness the results when following PAM or CLARA). This result suggests that SillyPutty’s unique approach of maximizing silhouette width can significantly enhance clustering outcomes compared to established methods.

The results displayed in Fig 4 also provide some insight into how some of the characteristics of a data set are likely to affect clustering results. The decrease in performance associated with more clusters (which provides more different ways to misclassify an individual sample) or more samples (which means more things that might be misclassified) or more noise were expected. The decrease in ARI and other measures in the presence of more features was unexpected. It may be explained by an increased difficulty of extracting the true signal in the presence of additional features that are entirely unrelated to the true subtype. This finding suggests that, with real data, it may be valuable to filter out features that do not seem to help separate clusters, such as those that do not vary much across the data set, before trying to cluster the data.

The promising performance of SillyPutty, especially when coupled with hierarchical clustering, opens up opportunities for its application in various domains. SillyPutty’s ability to enhance clustering outcomes could significantly impact fields like biology, where identifying distinct subgroups in large data sets is critical for understanding complex biological systems.

We cannot end this manuscript without mentioning some of the limitations of the present study. One of the current applications of clustering to biology lies in the realm of single cell RNA sequencing (scRNA). Such data sets frequently contain tens of thousands or even millions of individual cells that need to be clustered. Our simulations were limited to clustering 1000 samples. While that number is adequate for modern day studies of bulk samples, we cannot yet say that the performance will remain the same with much larger data sets. The speed of the best hybrid combination (hierarchical plus SillyPutty) suggest that it will be feasible to test that combination. But the time taken to run `SillyPutty` from 100 random starts would be a challenge. In principle, there are several ways to speed up the algorithm. For example, one could reduce the time to compute silhouette widths when moving one sample to another cluster by only recomputing values for the two clusters that changed. It might also be possible to recluster more than one sample during each iteration, though research would need to be done to determine what the optimal “move size” should be. In addition, it might be possible to transport some of the computations either into a faster compiled language or onto a parallel processing framework.

A second potential application area in biology or medicine is in the area of clinical data, which is often characterized as “mixed”, since it contains a mixture of binary data (symmetric or asymmetric), categorical data (nominal or ordinal), and continuous data on widely divergent scales. The challenge here is determining a reasonable distance metric. Again, we have not performed studies on this kind of data, so we can not assert that the best method found in our paper will remain the best in that context. However, the Umpire package that we developed previously [21] and used to generate the data sets here has already been extended to be able to simulate mixed clinical data [22] and shown to be useful [24].

### 4.1 Conclusion

In conclusion, `SillyPutty` demonstrates great promise as a valuable addition to the toolkit of clustering practitioners. It is competitive with the best existing clustering algorithms. Its effectiveness in improving clustering outcomes warrants continued investigation and exploration across a diverse range of data sets and real-world applications.

### 4.2 Availability

The `SillyPutty` R package has been submitted to the Comprehensive R Archive Network (CRAN).

Code to perform and analyze the simulations described here can be found in a Git project hosted at https://gitlab.com/krcoombes/sillyputty.

## Acknowledgments

The research described here was funded using startup funds from the Georgia Cancer Center at Augusta University and from an NIH grant, R25 MD011712, titled BD4ISU: Big Data for Indiana State University.

